# ELO-6 expression predicts longevity in isogenic populations of *Caenorhabditis elegans*

**DOI:** 10.1101/2024.03.23.585832

**Authors:** Weilin Kong, Guoli Gu, Tong Dai, Beibei Chen, Mintie Pu

**Affiliations:** State Key Laboratory of Conservation and Utilization of Bio-resources in Yunnan, Center for Life Sciences, School of Life Sciences, Yunnan University, Kunming, China

## Abstract

Variations of individual lifespans within genetically identical populations in homogenous environments are remarkable, with the cause largely unknown. Gene expression changes with age, and the transcriptome changes correlate with chronological aging. Here, we show that in *Caenorhabditis elegans*, the expression dynamic of the fatty acid elongase ELO-6 during aging predicts individual longevity in isogenic populations. The expression of *elo-6* is reduced with age. From adult day 5, ELO-6 expression level exhibits variation between individuals, and the expression level is positively correlated with adult lifespan and health span. Interventions that prolong longevity enhance the expression stability of ELO-6 during aging from adult day 4 to adult day 8, indicating ELO-6 is also a populational lifespan predictor. We performed transcriptome analysis in short-lived and long-lived isogenic worms and identified differentially expressed genes, which are enriched for PQM-1 binding sites. Decreasing *pqm-1* expression in young adults improved the homogeneity of ELO-6 levels between individuals and enhanced health span. Furthermore, we found reducing the expression of genes that are highly expressed in short-lived individuals, including PQM-1 target genes, enhanced ELO-6 expression stability with age and extended lifespan. Thus, our study identified ELO-6 as a predictor of individual and populational lifespan and revealed the role of *pqm-1* in restricting health span and possibly causing individual lifespan variation.

## Introduction

In laboratory animals and in humans, 60–90% of the variation in lifespan is independent of genotype ^1^. Genetically identical invertebrates in homogenous environments also experience similar variability in lifespan, indicating neither genetic nor environmental factors fully account for variability in individual longevity ^1^. Thus far, a single reporter gene that functions as a biomarker of individual aging, which will facilitate the investigation of stochastic factors causing individual lifespan variation, remains rare. Previous studies in *Caenorhabditis elegans* have found that *hsp-16.2* induction under heat shock, mitoflash frequency, nucleolus size, miRNA expression, and ROS level in the L2 developmental stage predict individual longevity, indicating *C. elegans* is a powerful model organism to investigate individual lifespan variation ^2–6^.

Transcriptome analysis in multiple organisms including yeast, nematodes, fruit flies, mice, and humans found that significant changes in gene expression occurs during aging ^7–9^. These age-regulated genes generally participate in a wide variety of biological processes that are associated with age-dependent physiological changes. In mice, most age-regulated gene expression alterations can either be completely or partially delayed by caloric restriction, a well-known intervention that retards aging ^10^. The extension of lifespan achieved by pharmacological intervention is accompanied by an enhancement of the overall transcriptional stability during aging ^11^. These results suggest that gene expression regulation plays an important and causal role in the aging process. Studies demonstrate that transcription profiles can serve as a cellular ageing clock for measuring age ^12–14^, indicating that gene transcriptional changes can be a potential tool to identify biomarkers of aging. PQM-1 is a C2H2 Zn finger transcription factor which binds to genes involved in lipid metabolism ^15^. PQM-1 binds to the DAF-16 associated element (DAE) and functions as an important mediator of the insulin signaling pathway. It is essential for the long-lived *daf-2* mutant phenotypes of somatic aging, stress response, and reproductive aging ^16,17^. PQM-1 is also essential for the lifespan extension of *puf-8* mutant animals ^18^. Despite its role in the lifespan extension of *C. elegans* mutant worms, *pqm-1* mutants themselves have a wild-type lifespan, and overexpressing PQM-1 protein shortens lifespan ^16^. Under stress conditions, PQM-1 limits survival from hypoxia and mediates the tradeoff between immunity and longevity ^19,20^.

In this study, we sought to identify lifespan predictors through transcriptome longitudinal analysis in *C. elegans* somatic cells during aging. We identified genes that continuously increased or decreased expression along with age, and further selected candidates by the correlation coefficient between gene expression abundance and age, the gene expression abundance in young adults, gene body histone H3.3 deposition, and histone modification patterns. We chose genes which showed a consistently reduced expression with age as candidates to be a lifespan predictor. ELO-6, a fatty acid elongase, had continuously decreased expression with age. From adult day 5, ELO-6 showed a variable expression level between individual animals, and the expression level predicted individual lifespan and health span. In addition, interventions prolonging lifespan enhance ELO-6 expression stability with age, indicating ELO-6 is a predictor of individual lifespan and populational lifespan. Through transcriptome analysis of the short-lived and long-lived animals distinguished by ELO-6 expression on adult day 5, we found that *pqm-1* regulates ELO-6 expression homogeneity in young adults and impacts health span. In addition, decreasing the expression of genes highly expressed in short-lived individuals, including genes targeted by PQM-1, extends lifespan. Therefore, our study identified a single gene reporter for individual and populational lifespan prediction and suggested the function of *pqm-1* in individual lifespan and health span regulation.

## Results

### Screen for lifespan predictors through transcriptome analysis

Gene expression profiles during aging are effective for age calculation ^12–14^. To identify genes that have the potential for lifespan prediction, we generated whole worm gene expression profiles of *C. elegans* somatic cells on adult day 2 (D2A), D4A, D6A, D8A, D10A, and D12A from germline-less *glp-1(e2141)* growing at 25 °C by mRNA-seq and then performed a longitudinal analysis. The MDS plot shows that the transcription profiles from young to old are arranged mainly from left to right on dimension 1 which represents 51% of the gene expression difference (Figure S1A). When analyzing the gene expression change at the single gene level, most gene expression fluctuated during aging (Table S2). We used 20% of FPKM (fragments per kilobase of transcript per million fragments mapped) as the cutoff for gene expression change between adjacent time points. We found 123 genes consistently increased expression and 44 genes consistently decreased expression during aging (Table S2). GO term analysis showed that the 44 genes that decreased expression with age are enriched for genes involved in innate immune response, structural constituent of cuticle, and oxidoreductase activity (Table S3), implicating physiological deterioration during aging. We aim to identify genes for lifespan prediction and further develop downstream screening assays. Therefore, we sought to identify lifespan predictors from the 44 genes that decreased expression with age. To select preferred candidates from the 44 genes, we first calculated the correlation coefficient between gene expression abundance and age (Table S2). Among the 44 genes, the expression level of *elo-6* is mostly inversely correlated with age, with a correlation coefficient of -0.98. In addition, *elo-6* is highly expressed on D2A compared with the other genes that have a similar inverse correlation coefficient between expression abundance and age (Table S2). Consistent with its active expression in young worms, *elo-6* is with HIS-72, one of the histone H3 variant H3.3 in *C. elegans*, deposited on the gene body region in somatic cells (Figure S2A)^21^. H3.3 is related to active gene expression ^22^ and is essential for the lifespan extension of long-lived mutants ^23^. We found a loss-of-function mutation of *his-72* shortens the lifespan of N2 (Figure S2B). Therefore, the expression of *elo-6* may be related to lifespan regulators. In addition, *elo-6* was found to possess an adult-specific histone modification pattern, with adult-specific H3K4me3 instead of H3K36me3 on its gene body region. (Figure S2A) ^21,24^. Therefore, we first examined whether *elo-6* can be a lifespan predictor.

We used the strain MTP27 with the endogenous ELO-6 tagged by GFP to examine the expression change of ELO-6 with age. This strain was generated by CRISPR/Cas9 by adding a GFP::3xFLAG tag to the N terminus of ELO-6. MTP27 develops normally and has a lifespan similar to N2 (Figure S3). GFP::ELO-6 is mainly located in the intestine, as previously reported (Figures 1A and 1C) ^25^. We observed that GFP::ELO-6 shows reduced expression during aging in N2 at 20 °C (Figures 1A and 1B) and *glp-1(e2141)* at 25 °C (Figures 1C and 1D). On D9A in N2 at 20 °C and D7A in *glp-1(e2141)* at 25 °C, the GFP signal can barely be observed under a fluorescence stereo microscope, which is consistent with the mRNA-seq result that the mRNA abundance of *elo-6* on D10A is 10% of that on D2A (Table S1). Thus, ELO-6 expression is an ideal candidate to monitor aging.

**Figure 1.**
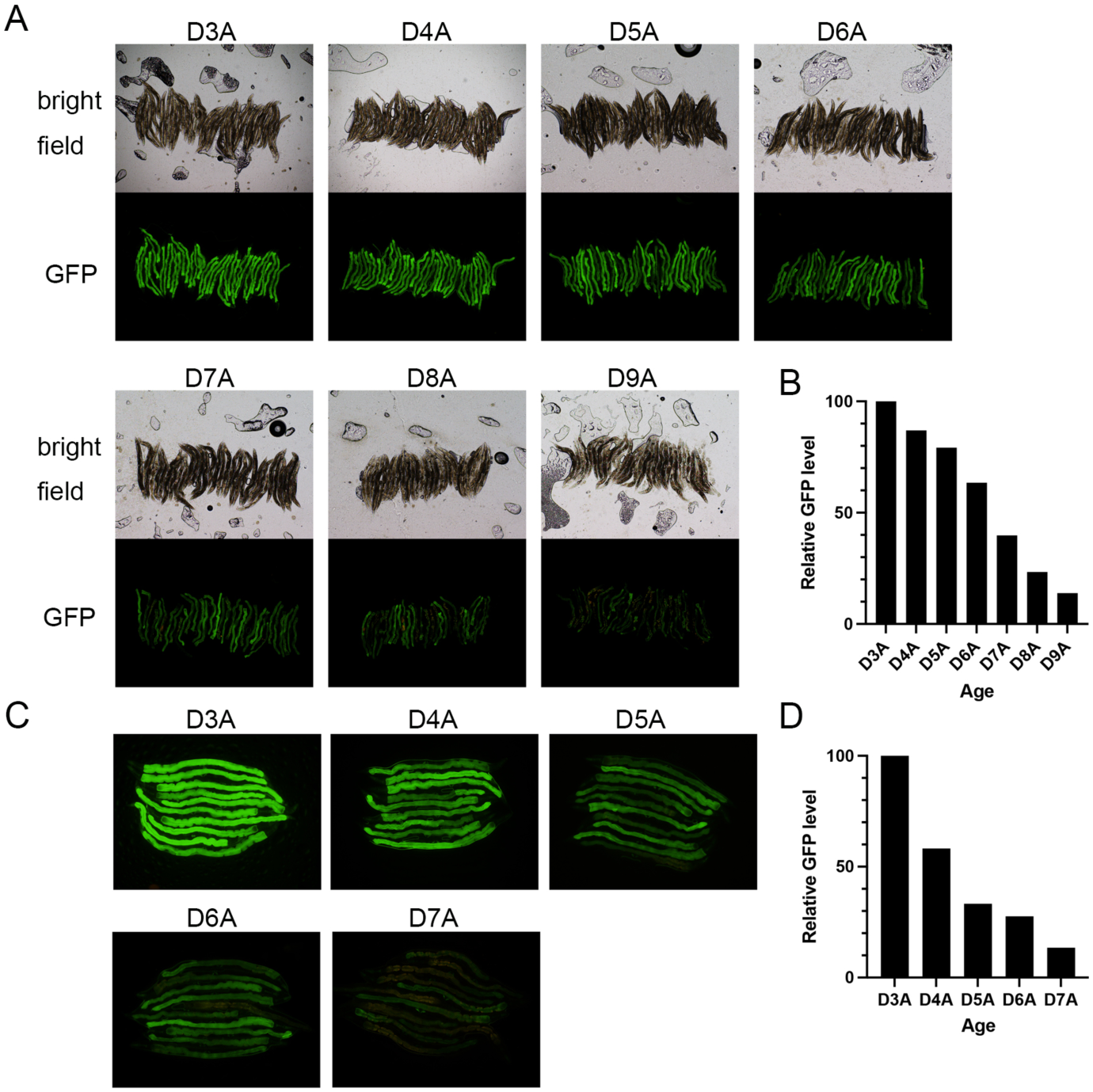
The expression level of GFP::ELO-6 in the intestine is consistently reduced during aging. (A) The expression of GFP::ELO-6 in intestine in MTP27*(gfp::3xflag::elo-6)* is consistently reduced from D3A to D9A. Images were taken with a 5x objective lens with consistent imaging parameter settings. (B) The total GFP signal in each image in (A) was quantified by ImageJ. The mean GFP level *per* worm is presented. (C) The expression of GFP::ELO-6 in intestine in MTP29*(gfp::3xflag::elo-6;glp(e2141))* is consistently reduced from D3A to D7A. Images were taken with a 10x objective lens with consistent imaging parameter settings. (D) The total GFP signal in each image in (C) was quantified by ImageJ. The mean GFP level *per* worm is presented.

### Expression variation of ELO-6 in young adults predicts lifespan and health span in isogenic populations

We observed that as early as D5A, there were differences in GFP::ELO-6 expression levels between individual worms in the N2 background (Figure 1A) and on D4A in *glp-1(e2141)* (Figure 1C). Since ELO-6 expression decreases with age, we hypothesized that the variation of ELO-6 expression represents the difference in the aging process between individuals. We separated worms according to GFP::ELO-6 levels on D5A in N2 background into 3 groups (Figure 2A) and measured their lifespan (Figure 2B, Table S8). The worms with low GFP::ELO-6 expression (GFPL) have a significantly shorter lifespan than those of medium (GFPM) and high GFP::ELO-6 (GFPH) expression groups (15.23% and 17.20% mean lifespan difference, *P-*value < 0.01, *P*-value < 0.01)(Figure 2B, Table S8), while GFPM and GFPH have a similar lifespan (*P*-value = 0.815) (Table S8). In addition, we also measured the pharyngeal pumping rate and body bending rate, which represents the health span. Compared with GFPM and GFPH groups, the GFPL group show a significantly slower bending rate and pharyngeal pumping rate (Figures 2C and 2D), indicating that individuals with a more rapid reduction of ELO-6 expression during aging in the GFPL group also have shorter health spans. As well, the GFPH group shows a significantly higher pharyngeal pumping rate than the GFPM group animals on D8A (Figure 2D), indicating its relatively slower aging process. The offspring of GFPL, GFPM, and GFPH shows similar GFP expression on D5A (Figure S4C), confirming that the expression variation of ELO-6 on D5A is not caused by genetic factors.

**Figure 2.**
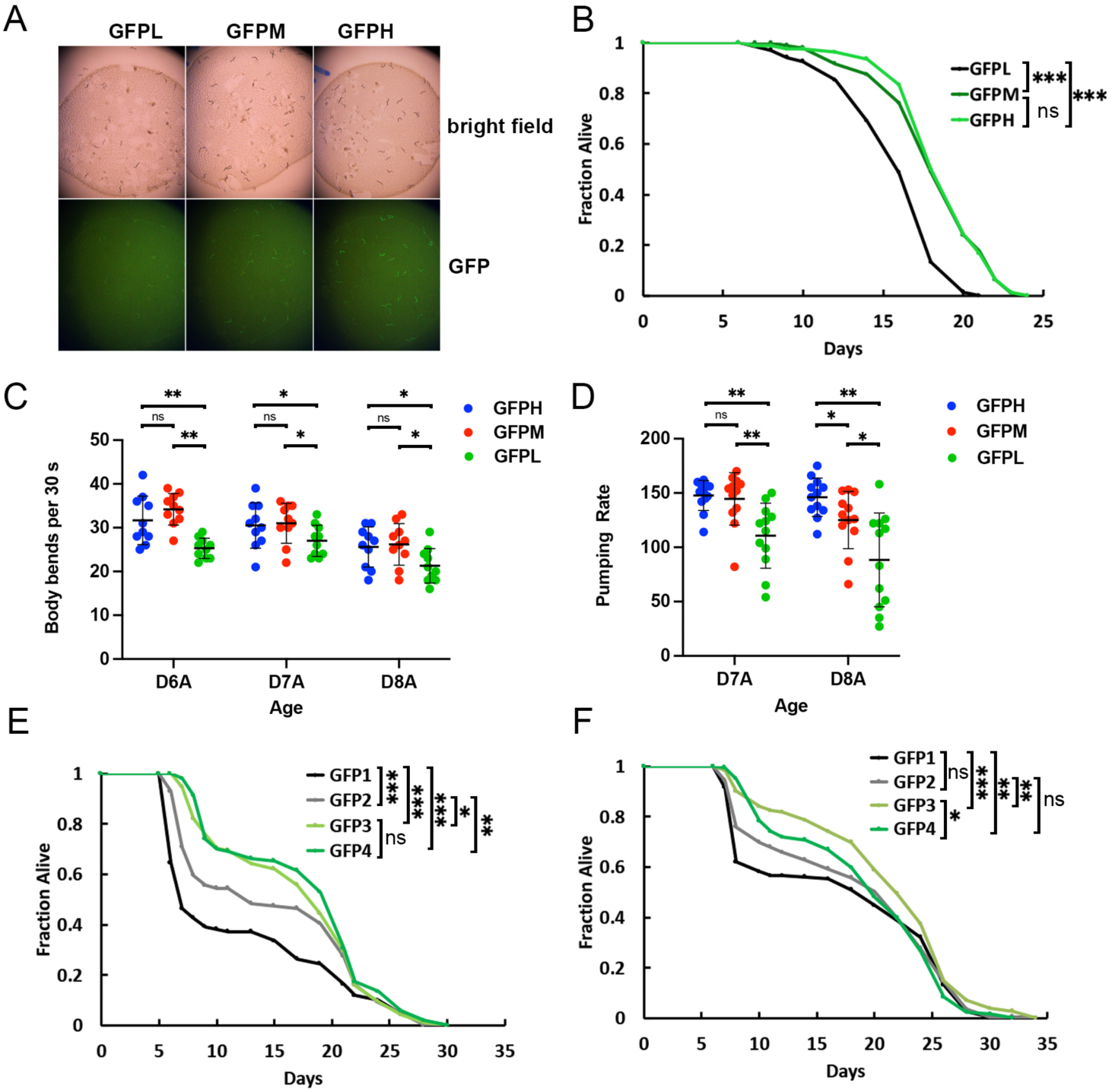
Expression variation of ELO-6 in young adults predicts lifespan and health span in isogenic populations. (A) The GFP expressions in GFPL, GFPM, and GFPH group worms from MTP27. Images were taken by a fluorescence stereo microscope with consistent imaging parameter settings. (B) GFPL animals have a shorter lifespan compared with GFPM and GFPH. Survival curves of GFPL, GFPM, and GFPH are shown. Quantitative data are presented in Supplementary Table S8. (∗∗∗) *P*-value < 0.001, *Log-rank test* (C) GFPL animals have a slower body bending rate compared with GFPM and GFPH on adult day 6, day 7, and day 8. (∗∗) *P*-value < 0.01, (∗) *P*-value < 0.05, (ns) *P*-value > 0.05, *unpaired t-test, two-sided except for GFPH vs GFPL on D7A only one-sided test showing significant difference*. (D) GFPL animals have a slower pharyngeal pumping rate compared with GFPM and GFPH on adult day 7 and day 8. GFPH animals have a faster pharyngeal pumping rate compared with GFPM on adult day 8. (∗∗) *P*-value < 0.01, (∗) *P*-value < 0.05, (ns) *P*-value > 0.05, *unpaired t-test, two-sided* (E) Animals with low GFP::ELO-6 expression on adult day 5 are short-lived in the *glp-4(bn2)* background. Survival curves of GFP1, GFP2, GFP3, and GFP4 are shown. The corresponding images are in Supplementary Figure S4A. Quantitative data are presented in Supplementary Table S8. (∗∗∗) *P*-value < 0.001, (∗∗) *P*-value < 0.01, (∗) *P*-value < 0.05, (ns) *P*-value > 0.05, *Breslow test* (F) Animals with low GFP::ELO-6 expression on adult day 5 are short-lived in the *glp-1(e2141)* background. Survival curves of GFP1, GFP2, GFP3, and GFP4 are shown. The corresponding images are in Supplementary Figure S4B. Quantitative data are presented in Supplementary Table S8. (∗∗∗) *P*-value < 0.001, (∗∗) *P*-value < 0.01, (∗) *P*-value < 0.05, (ns) *P*-value > 0.05, *Breslow test*

In germline-less *glp-4(bn2)* and *glp-1(e2141)* worms cultured at 25°C, we separated worms according to GFP levels on D5A from low to high into GFP1 to GFP4 groups (Figure S4A and S4B). We found that worms with lower GFP::ELO-6 expression at D5A had a shorter lifespan when analyzed using the Breslow test which is suitable for early-stage aging comparison (Figures 2E and 2F, Table S8). Thus, the expression level of ELO-6 on D5A is positively correlated with individual lifespan and health span in isogenic populations.

### ELO-6 is a populational lifespan predictor

We further asked whether ELO-6 can predict lifespan regulated by genetic factors, in addition to the non-genetic ones that influence individual lifespan. To answer this question we applied RNAi from hatching to reduce the expression of *daf-2*, *cyc-1*, *set-2*, and *utx-1*, which all extend lifespan ^26–30^, and examined GFP::ELO-6 expression during aging. *daf-2* RNAi enhanced ELO-6 expression stability as early as D4A, and showed a higher GFP level than the control on D7A (Figures 3A and 3B). This effect depends on *daf-16*. In the short-lived *daf-16(mu86)* mutant, the expression level of ELO-6 showed no difference between *daf-2* RNAi and the control (Figures 3A and 3B). In addition, ELO-6 expression on D6A in *daf-16(mu86)* under control treatment is reduced compared with the N2 worms (Figures 3A and 3B). Under *utx-1* RNAi treatment, ELO-6 expression stability was enhanced during aging (Figures 3C and 3D). GFP::ELO-6 expression can still be observed on D9A when the GFP expression in the control condition can barely be detected (Figure 3C). Under *cyc-1* RNAi treatment, the expression level of GFP::ELO-6 is lower than that of the control from the L4 stage (data not shown) and shows a slower decrease during aging from D2A to D8A (Figures 3C and 3D), indicating ELO-6 expression stability is enhanced by *cyc-1* RNAi as well. *Set-2* RNAi extends lifespan (Figure S5A, Table S8). The ELO-6 expression level is only increased by *set-2* RNAi on D4A, and the expression decrease of ELO-6 is similar to the control from D6A to D8A (Figures 3A and 3B). Germline-less animals have prolonged longevity ^31^. We further tested whether ELO-6 expression can also be stabilized during aging in a long-lived germline-less mutant. We observed that in long-lived *glp-1(e2141)*, after being shifted from 25°C to 20°C after D1A ^32^, the ELO-6 expression was stabilized during aging (Figure 3E). Therefore, ELO-6 is a predictor of lifespan regulated by genetic factors. In addition, metformin treatment, which extends lifespan ^33^, also enhances the GFP::ELO-6 expression stability during aging (Figures 3F and 3G). These results demonstrate that the expression stability of ELO-6 during aging is a general predictor of populational lifespan.

**Figure 3.**
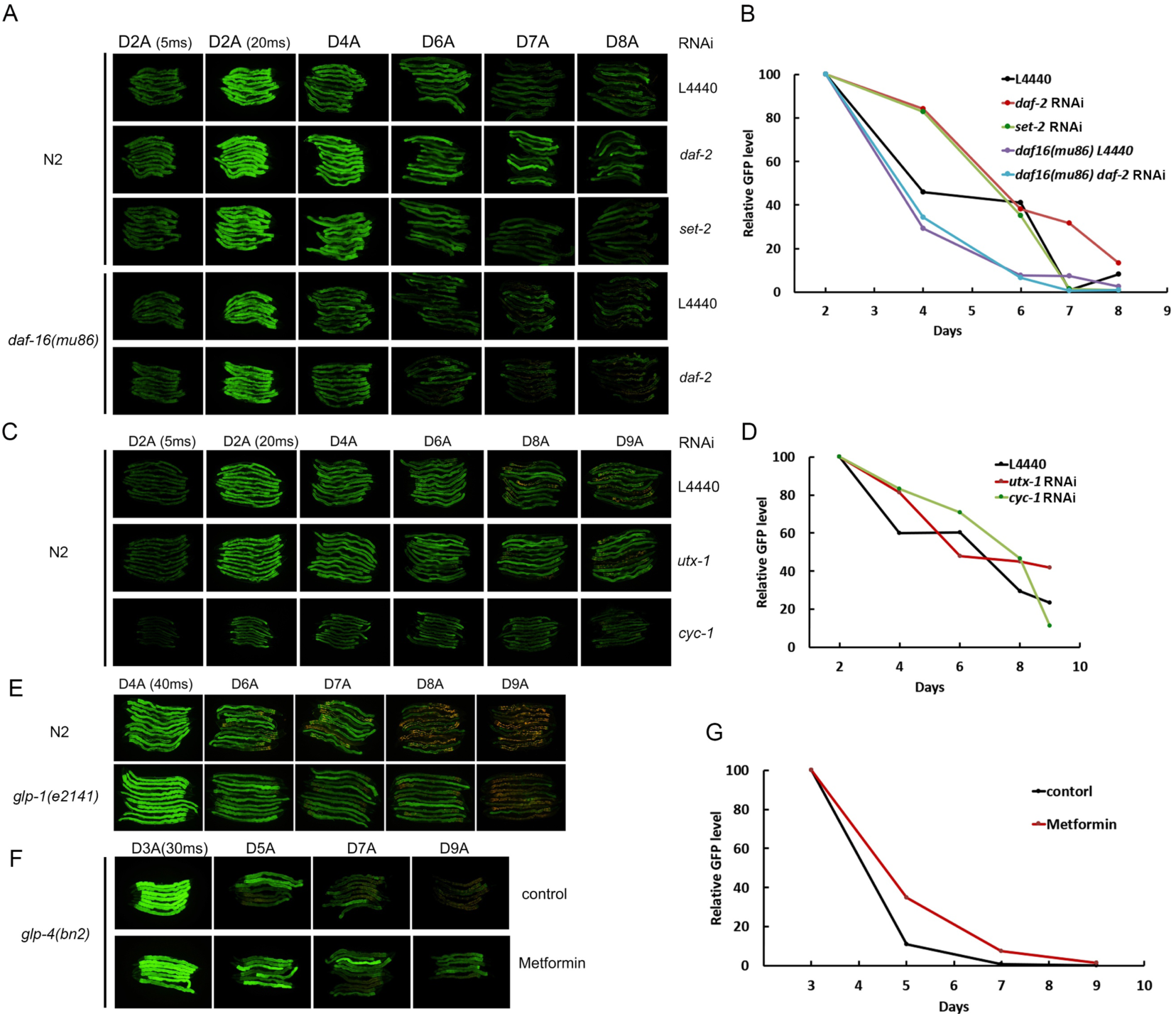
ELO-6 is a predictor of population lifespans regulated by genetic factors and metformin treatment. (A) *daf-2* RNAi treatment enhances ELO-6 expression stability from D4A. This increase of ELO-6 expression stability depends on *daf-16*. In the short-lived *daf-16(mu86)* strain, the expression stability of ELO-6 is decreased with age from D6A. *Set-2* RNAi enhances ELO-4 expression stability till D4A. For D2A worms, images taken with acquisition times of 5 ms and 20 ms are shown. For the other ages, images taken at 20 ms are shown. (B) The total GFP signal in each image in (A) was quantified by ImageJ with color threshold hue setting 50-255 to avoid the autofluorescence signal. The mean GFP signal *per* worm is presented. (C) *utx-1* and *cyc-1* RNAi treatment enhance ELO-6 expression stability with age. For D2A, images taken with acquisition times of 5 ms and 20 ms are shown. For the other ages, images taken at 20 ms are shown. (D) The total GFP signal in each image in (C) was quantified by ImageJ with color threshold hue setting 50-255 to avoid the autofluorescence signal. The mean GFP signal *per* worm is presented. (E) ELO-6 expression stability is increased in long-lived germline-less mutant *glp-1(e2141)*. (F) Metformin treatment enhances ELO-6 expression stability with age. (G) The total GFP signal in each image in (F) was quantified by ImageJ with color threshold hue setting 50-255 to avoid the autofluorescence signal. The mean GFP signal *per* worm is presented.

Since the ELO-6 expression during aging is correlated with lifespan, we further asked whether *elo-6* is a lifespan regulator. We reduced *elo-6* expression by RNAi from D2A or D5A, and found that *elo-6* RNAi does not change lifespan (RNAi from D2A, *P*-value = 0.166; RNAi from D5A, *P*-value = 0.667) (Figures S6A and S6B, Table S8). Supplementation of C17iso, the product of ELO-6 ^25^, can only slightly extend the populational lifespan when the treatment begins at D4A (3% mean lifespan extension, *P*-value = 0.03) (Figure S6C, Table S8) and does not affect the lifespan of the GFPL group when the treatment is started on D6A (*P*-value = 0.851) (Figure S6D, Table S8). Therefore, *elo-6* expression in the intestine is not a lifespan regulator but simply a lifespan predictor. Since ELO-6 expression stability during aging is influenced by the insulin/IGF-1 signaling pathway, and previous studies showed that the *daf-16* downstream gene *sod-3* shows expression variation between individuals on D9A reflecting the pathogenic effect of OP50 ^34^, we asked whether the expression variation of ELO-6 on D5A can be caused by OP50 infection as well. We transferred MTP27 for 2 generations onto OP50 plates treated with UV, and found that GFP expression on D5A still shows individual variation (Figure S7), indicating that ELO-6 expression variation on D5A is not caused by OP50 infection.

### Transcriptome analysis of short-lived and long-lived individuals

To understand the molecular mechanism that is related to ELO-6 prediction of individual worm lifespan, we performed mRNA-seq to identify differentially expressed genes (DEGs) between the short-lived and long-lived animals distinguished by ELO-6 expression variation on D5A. We performed mRNA-seq in both germline-less mutant *glp-1(e2141)* and *glp-4(bn2)* backgrounds to avoid the bias caused by *glp-1(e2141)* or *glp-4(bn2)* mutation. On D5A, we separated the isogenic worms into GFP1, GFP2, GFP3, and GFP4 groups according to GFP levels from low to high, similar to the grouping for lifespan analysis, and pooled 100 worms in each group for mRNA-seq. Transcriptome analysis showed no striking gene expression differences between different GFP groups, with all correlation coefficients higher than 0.85 (Table S1). The MDS plot of mRNA-seq data showed that the difference of transcriptomes followed similar trends as the GFP signal intensity (Figure 4A). Comparing mRNA-seq data from adjacent GFP groups from the *glp-1(e2141)* background showed that GFP1 vs GFP2 groups had 73 significantly differentially expressed genes when applying FDR <0.1, while there were only 40 genes between GFP2 and GFP3, and 10 genes between GFP3 and GFP4 that showed expression differences (Table S4). Since we observed that there is a large fraction of the population that dies on D6A in GFP1 groups in both the *glp-1(e2141)* and *glp-4(bn2)* background (Figures 2E and 2F), the transcriptome of GFP1 might reflect the features that occur before death. Considering that the transcriptomes of GFP3 and GFP4 are highly similar, we then compared the transcriptome between GFP2 and GFP4 to identify differentially expressed genes (DEGs) between the shorter-lived and longer-lived groups. We found 65 genes increased expression and 113 genes decreased expression in the short-lived worms compared with the long-lived ones in both *glp-1(e2141)* and *glp-4(bn2)* backgrounds (Figure 4B). Functional clustering analysis showed that xenobiotic metabolic process and structural constituent of cuticle-related genes are enriched in the decreased expression genes, and ribosome biogenesis and CUB domain related genes are enriched in increased expression genes in short-lived worms (Table S5).

**Figure 4.**
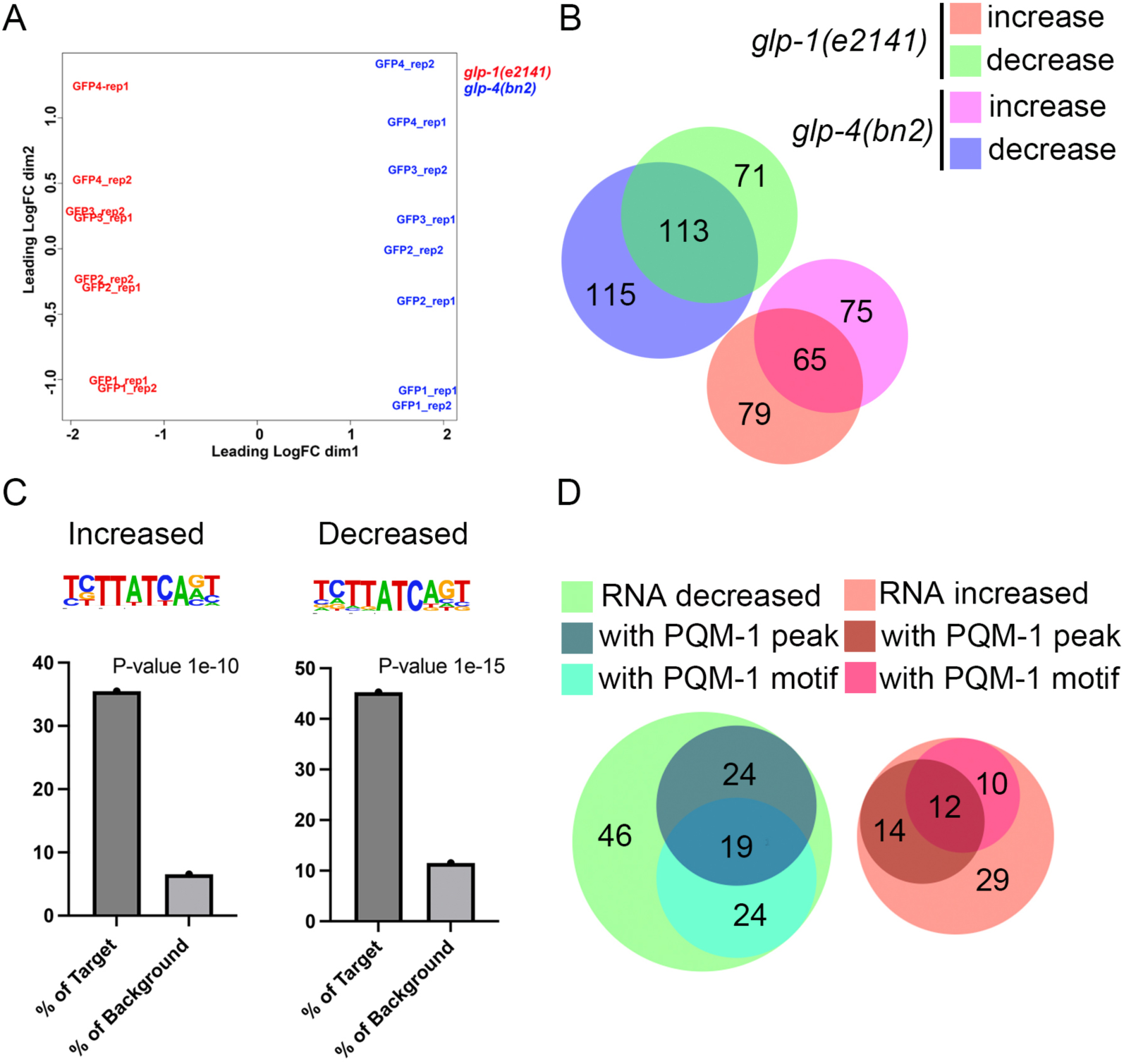
Transcriptome analysis of short-lived and long-lived individuals. (A) MDS plot of mRNA-seq performed in GFP1, GFP2, GFP3, and GFP4 animals in the *glp-1(e2141)* and *glp-4(bn2)* background. (B) The Venn diagrams show genes that were differentially expressed between GFP2 and GFP4 worms in the *glp-1(e2141)* and *glp-4(bn2)* background. Red: genes highly expressed (increased) in GFP2 in *glp-1(e2141)*; Lime: genes low expressed (decreased) in GFP2 in *glp-1(e2141)*; Fuchsia: genes highly expressed in GFP2 in *glp-4(bn2)*; Aqua: genes low expressed in GFP2 in *glp-4(bn2)*. Gene numbers for each group are shown. (C) Motif analysis with genes differentially expressed between GFP2 and GFP4 worms in both the *glp-1(e2141)* and *glp-4(bn2)* background. Genomic regions surrounding TSS [-400, +100] of the 65 increased genes and 113 decreased genes in GFP2 were analyzed by Homer. Genes that are highly expressed and low expressed in GFP2 animals are both enriched in the PQM-1/ELT-3 binding motif. TSS: transcriptional start site. (D) The Venn diagrams show the occupancy of PQM-1/ELT-3 binding motif and PQM-1 peak in differentially expressed genes between GFP2 and GFP4. The genes showing expression difference between GFP2 and GFP in both the *glp-1(e2141)* and *glp-4(bn2)* background were subjected to analysis. Gene numbers for each group are shown.

We further asked whether the DEGs between short-lived and long-lived isogenic animals are under a common regulatory mechanism. Motif analysis with the 65 increasingly expressed and 113 decreased expressed genes showed that both gene groups are enriched for the PQM-1/ELT-3 binding motif in the gene promoter regions (Figure 4C), indicating common transcription factors control the differentially expressed genes. In addition to the motif analysis, we inspected the PQM-1 ChIP-seq profile from L3 stage worms ^15^. In the 65 genes with increased expression, 26 have PQM-1 peaks in gene upstream regions, with 12 overlapping with the PQM-1/ELT-3 binding motif prediction. Of the 113 genes with decreased expression, 43 have PQM-1 peaks in gene upstream regions, and 19 coincided with PQM-1/ELT-3 binding motif prediction (Figure 4D, Table S6). Therefore, the transcriptome difference between short-lived and long-lived individuals might be regulated by PQM-1/ELT-3.

### *pqm-1* promotes ELO-6 expression variation between individuals in mid-aged adults and regulates health span

To investigate whether PQM-1 or ELT-3 regulates ELO-6 expression variation between individual worms on D5A and during subsequent aging, we used RNAi to knockdown *pqm-1* or *elt-3* starting from D2A or D3A and examined GFP::ELO-6 expression. We found that *pqm-1* RNAi decreased the variation of ELO-6 expression between individuals, with fewer worms with low GFP::ELO-6 expression on D6A compared to the control (Figures 5A and S8), and the phenotype is stronger than *elt-3* RNAi (data not shown). Therefore, we focused on *pqm-1* for further analysis. To test whether *pqm-1* regulates the aging process, we measured the worm bending frequency and pharyngeal pumping rate of the mid-aged worms upon *pqm-1* RNAi treatment starting from D2A. The results show that decreased *pqm-1* expression delayed the aging process, with the bending frequency on D6A and D8A being significantly higher in the *pqm-1* RNAi group than in the control group (Figure 5B). The pharyngeal pumping rate under *pqm-1* RNAi treatment is also higher than the control on D7A and day D9A (Figure 5C).

**Figure 5.**
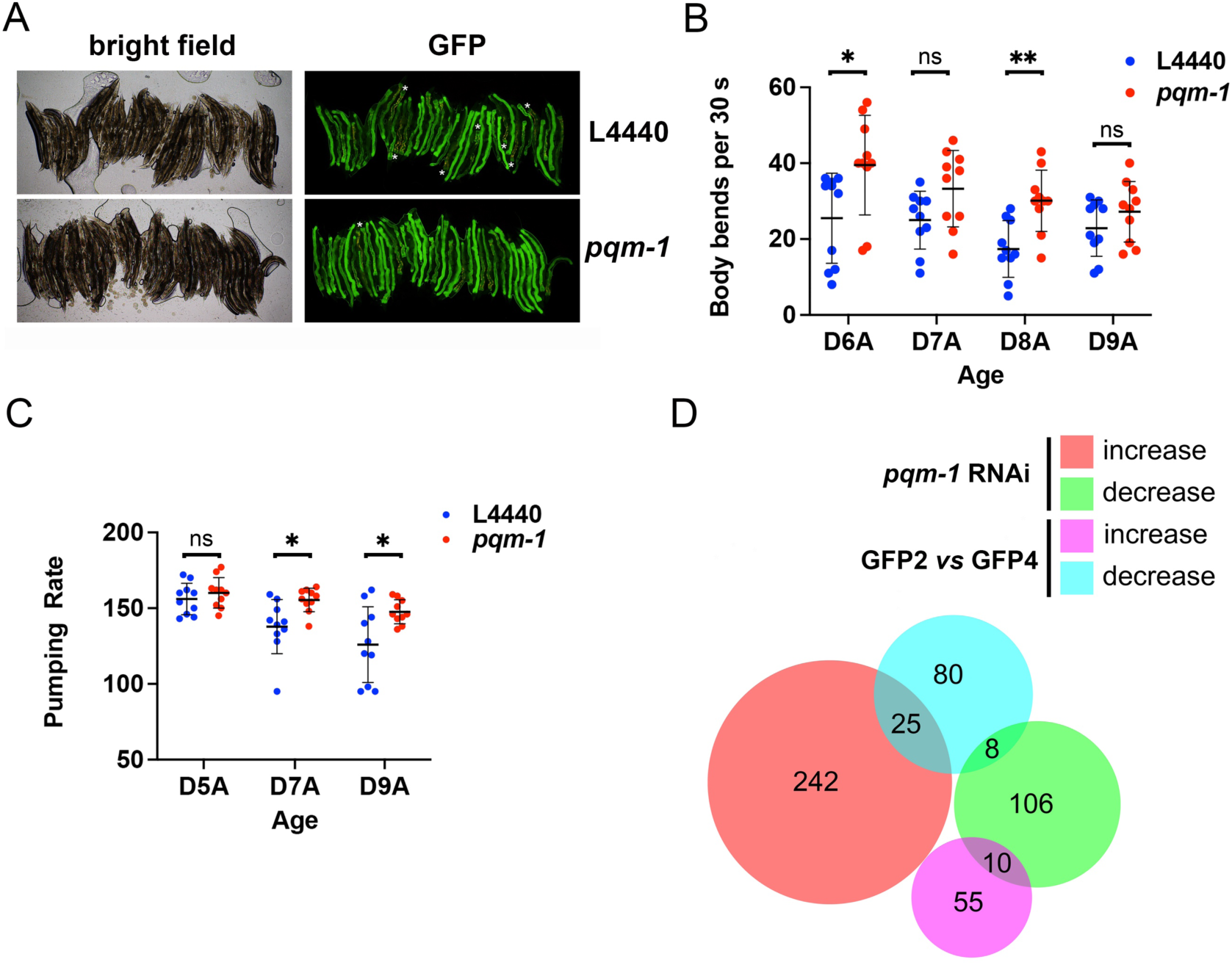
*pqm-1* promotes ELO-6 expression variation between individuals in mid-aged adults and regulates health-span. (A) *pqm-1* RNAi treatment enhances GFP::ELO-6 expression homogeneity in MTP27 on D6A. RNAi treatment started from D3A. *, worms without GFP signal in part of the intestine. 38 animals were in both RNAi groups. 7 worms or 1 worm are without GFP signal in part of the intestine under control or *pqm-1* RNAi treatment. The yellow signal is autofluorescence. Additional replicates are shown in Supplementary Figure S8. (B) *pqm-1* RNAi animals have a faster body bending rate on D6A and D8A. RNAi treatment started from D2A. (∗∗) *P*-value < 0.01, (∗) *P*-value < 0.05, *unpaired t-test, two-sided* (C) *pqm-1* RNAi animals have a faster pharyngeal pumping rate on D7A and D9A. RNAi treatment started from D2A. (∗∗) *P*-value < 0.01, (∗) *P*-value < 0.05, *unpaired t-test, two-sided* (D) Differentially expressed genes caused by *pqm-1* RNAi treatment and those between GFP2 and GFP4 are compared with Venn diagrams. Red: genes increased expressed (increased) upon *pqm-1* RNAi treatment; Lime: genes with reduced expression (decreased) upon *pqm-1* RNAi treatment; Fuchsia: genes highly expressed in GFP2; Aqua: genes low expressed in GFP2. Gene numbers for each group are shown.

We further examined the transcriptome change on D5A upon *pqm-1* RNAi treatment starting from D2A in the germline-less mutant *glp-1(e2141)* growing at 25 °C. The MDS plot of the mRNA-seq data from the *pqm-1* RNAi treatment along with the transcriptome data from D2A to D12A shows that the transcriptome upon *pqm-1* RNAi is shifted towards younger age on dimension 1 which represent 38% gene expression difference (Figure S1B). There were 267 genes with increased expression upon *pqm-1* RNAi treatment (fold change > 0.5, FDR < 0.05) and 124 genes with reduced expression (fold change > 0.5, FDR < 0.05) compared to the control. We then further compared the DEGs between the short-lived and long-lived individual animals (Figure 4B) with the expression difference caused by *pqm-1* RNAi treatment. We found that 10 out of the 65 genes that are highly expressed in short-lived worms had decreased expression upon *pqm-1* RNAi treatment, and among the 113 genes that are expressed at a lower level in short-lived worms, *pqm-1* RNAi treatment increased the expression of 25 genes and reduced the expression of 8 genes (Figure 5D). Thus, *pqm-1* RNAi influences the transcriptome to reduce the transcriptional difference that would occur between short-lived and long-lived individuals during aging.

### Reduction of the expression of genes highly expressed in short-lived individuals enhanced ELO-6 expression stability and regulated lifespan

We next investigated whether the DEGs between the short-lived and long-lived worms at D5A regulate GFP::ELO-6 expression stability during aging and lifespan. In this study, we focused on the genes with increased expression in the short-lived GFP2 group. We knocked down 61 of the 65 genes by RNAi treatment from D2A and examined the GFP level change with age and lifespan in the N2 background. We found that RNAi of 18 genes delayed the decrease of GFP::ELO-6 expression with age (Figures 6A-E, S9, Table S7), with 10 out of the 18 genes possessing a PQM-1 binding motif and/or PQM-1 peaks in the gene upstream regions (Table S7). Interestingly, among the 18 genes, *smf-2* and *B0238.13* reduces expression upon *pqm-1* RNAi treatment (Table S6), although there is no PQM-1 binding motif or PQM-1 binding site identified by ChIP-seq in their gene upstream region. Among the 18 genes, RNAi of ribosomal genes *tag-151, E02H1.1*, *C16A3.6*, *B0511.6*, and *T23D8.3* extended lifespan in the N2 background (Figure 6F, Table S8). RNAi *tag-151* enhanced the GFP signal on D6A (Figure 6A), and RNAi of *C16A3.6*, *E02H1.1*, *B0511.6*, *T23D8.3* delayed GFP level reduction until D8A (Figures 6B and 6C). The additional 3 genes, *nst-1*, *F53F4.11*, and *lpd-7*, when knocked down through RNAi, also extended the lifespan, while not having an obvious phenotype on the ELO-6 expression stability during aging (data not shown). Interestingly, all eight of these lifespan-regulating genes are related to nucleolus function (Table S7), which is consistent with previous findings that long-lived mutants are with reduced ribosomal function and small nucleoli are correlated with extended longevity in multiple organisms ^4,35^. Among these 8 genes, 3 of them have PQM-1 binding motif and/or PQM-1 peaks in the gene upstream regions (Table S7), including *T23D8.3*, which extends lifespan most profoundly (Table S7). *T23D8.3* has been identified as a longevity regulator through longevity network prediction ^36^. Therefore, ELO-6 expression and lifespan are regulated by these genes highly expressed in the short-lived animals. Understanding the regulatory mechanism that influences the expression of these genes will elucidate the cues that trigger the difference in the aging process between individuals.

**Figure 6.**
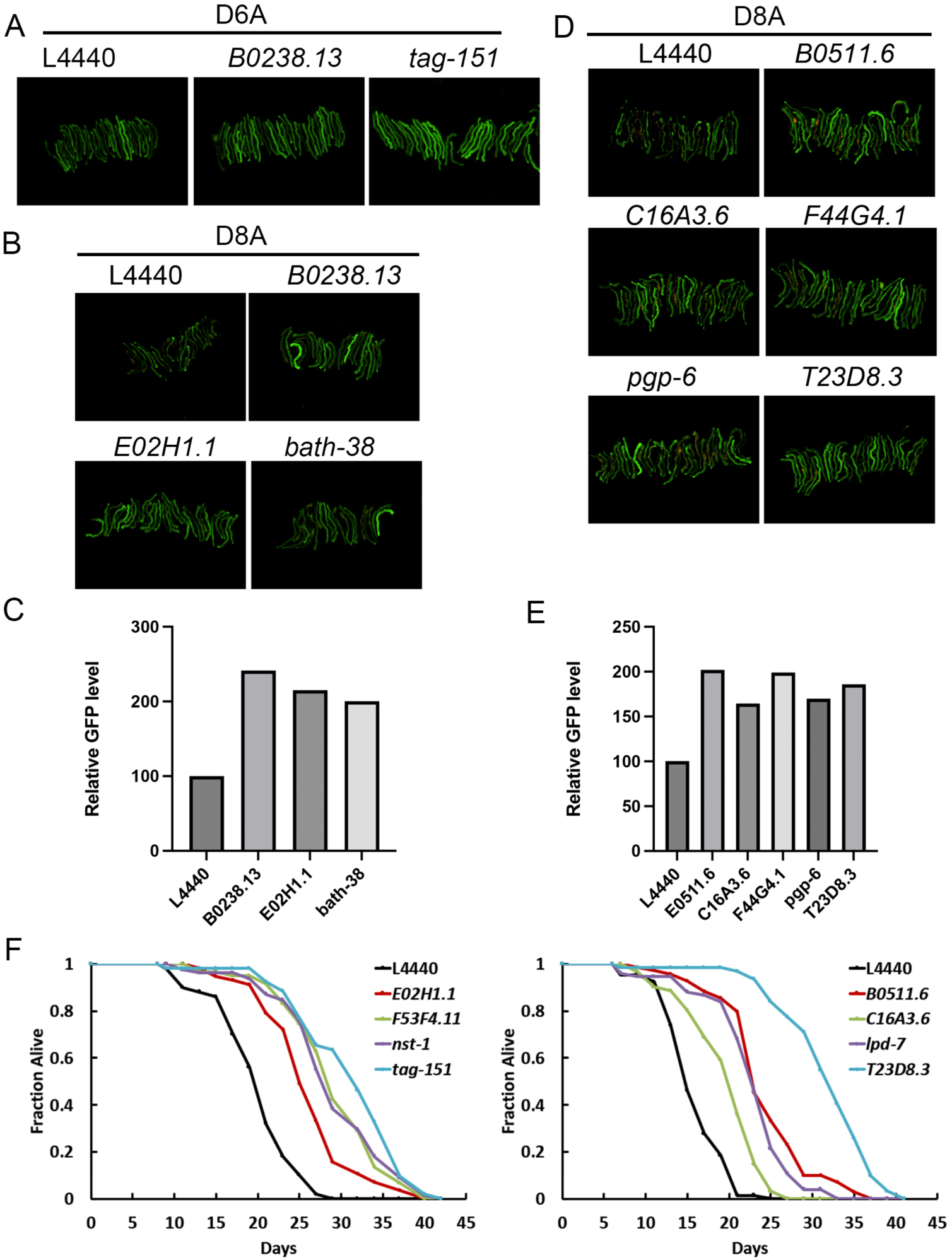
RNAi treatment of genes highly expressed in short-lived individuals enhance ELO-6 expression stability and extend lifespan. (A) RNAi treatment of *tag-151* and *B0238.13* from D2A enhances ELO-6 expression homogeneity on D6A. (B and D) Representative images showing RNAi treatment of the genes highly expressed in short-lived individuals from D2A enhances ELO-6 expression till D8A. (C and E) The total GFP signal in each image in (B) and (D) was quantified by ImageJ with color threshold hue setting 50-255 to avoid the autofluorescence signal. The mean GFP signal *per* worm is presented. (F)Reducing the expression of ribosome-related genes that are highly expressed in short-lived individuals extends lifespan. Quantitative data are presented in Supplementary Table S8.

## Discussion

In this study, we used longitudinal analysis of transcriptome profiles during aging to identify lifespan predictor genes. Generally, the transcriptome changes remarkably with age. In this study, when analyzing the transcriptome at an interval every 2 days during aging, we found only a few genes that could be identified as significantly changed in expression using the general RNA-seq data analysis pipeline. When combining the data from all aging intervals to identify genes with consistent trends of expression change, there were no genes which fulfilled the criteria. This suggests that gene expression during aging occurs with subtle changes. In this study, we used an arbitrary 20% FPKM difference as the criteria for gene expression changes between adjacent aging time points. We found 123 genes consistently increased and 44 genes consistently decreased expression with age. When inspecting the expression of the lifespan predictor candidate gene *elo-6*, we found that ELO-6 expression decreased consistently, as expected, indicating the arbitrary criteria is meaningful to measure trends of gene expression change during aging.

We determined that *elo-6* reduces expression from D4A according to mRNA-seq and GFP signal levels, and that it showed variation in expression between individual worms on D5A. This difference between individuals is neither caused by genetic variations nor by OP50 infection. We speculate that gene expression variation between individual animals during aging is not unique to *elo-6*, but should be a common feature for dynamically expressed genes with age. Other genes with expression differences between long-lived and short-lived individuals on D5A and also consistently changed expression levels during aging might be proper reporters as well (Table S2). The reasons why *elo-6* shows differences between individuals may be that *elo-6* is highly expressed in young adults with an expression level ranked 224 out of all the expressed genes on D2A, and *elo-6* expression is dramatically reduced during subsequent aging. Therefore, the abundance of expression change is adequate for detecting differences between individuals.

In the N2 background, the GFPL animals are significantly short-lived compared with the GFPM and GFPH group animals. *elo-6* encodes a fatty acid elongase synthesizing C17iso, and it is partially functionally redundant with *elo-5* ^25^. Unlike *elo-5*, which is essential for development, *elo-6* deficiency does not cause developmental defect ^25^. While ELO-6 expression from D5A is positively correlated with lifespan and health span, we found that RNAi of *elo-6* in adult stages does not affect lifespan. A previous study showed that C17iso supplementation from D1A does not affect lifespan ^37^. Therefore, we supplied C17iso from D4A and found that the supplement slightly extended the whole populational lifespan, which might reflect the lifespan extension of the short-lived animals. Therefore, we supplied C17iso to the GFPL group from D6A, but could not extend the lifespan of the GFPL group animals.

We also found that the expression dynamics of *elo-6* is a populational lifespan predictor. In this study, reducing the expression of *daf-2* and *utx-1*, which extends lifespan through the insulin/IGF-1 signaling pathway, enhances the expression stability of *elo-6* during aging. Accordingly, in the short-lived *daf-16(mus86)* mutant, the ELO-6 expression level was reduced earlier than wild-type N2. In the interventions extending lifespan we tested in this study, *set-2* RNAi does not change the ELO-6 expression stability with age after D6A, nor the ELO-6 expression level on D2A, indicating neither the active expression of *elo-6* in young animals, nor the expression stability during aging is controlled by *set-2*. The ELO-6 expression stability during aging is also enhanced in germline-less mutants and with metformin treatment. Therefore, ELO-6 expression dynamics can predict longevity under diverse interventions.

To investigate the mechanism in regulating individual lifespan, we performed mRNA-seq to compare the transcriptome of short-lived and long-lived isogenic worms distinguished by ELO-6 expression in germline-less mutants background. Surprisingly, the genome transcriptomes show high similarity between different GFP groups, despite the GFP1 groups with many worms dying the next day in both germline-less mutants. We were still able to identify genes with significantly changed expression in short-lived animals compared to long-lived ones. Xenobiotic metabolic process and structural constituent of cuticle-related genes are enriched in downregulated genes in short-lived worms, and ribosome biogenesis and CUB domain related genes were enriched in upregulated genes in short-lived worms. The PQM-1 binding site was found to be enriched in both downregulated and upregulated genes. Upon *pqm-1* RNAi treatment, transcriptome of the animals shifted to a relatively younger state, and the health span was improved. This is consistent with the previous finding that overexpression of PQM-1 shortens wild-type *C. elegans* lifespan ^16^. Since PQM-1 mediates the tradeoff between survival and lifespan under stress conditions, we speculate that the short-lived individuals might undergo certain stress conditions, which might be associated with malfunction of xenobiotic metabolic process, cuticle properties, or enhanced expression of ribosomal synthesis genes. In this study, we found that ribosomal genes are enrich in the highly expressed genes in short-lived animals, and RNAi of some of them extended lifespan. This is consistent with the previous findings that small nucleoli on adult day 1 predicts longer individual lifespan ^4^. C17iso mediates the coordination between growth and amino acids supplement under a dietary restriction condition ^37^. Enhanced expression of ribosomal synthesis genes in short-lived animals might reflect enhanced ribosomal function and, therefore, cause amino acids supplement stress, leading to the downregulation of *elo-6* to reduce the synthesis of C17iso.

Our study identified *elo-6* as a single gene reporter for prediction of individual and populational lifespan. By investigating the mechanism PQM-1 regulates the differentially expressed genes between short-lived and long-lived isogenic animals will help to understand the lifespan variation between individuals.

## Methods

### *C. elegans* strains and culture

*C. elegans* strain grown under standard growth conditions ^38^. All strains were grown and maintained on NGM plates seeded with *Escherichia coli* OP50 except for the RNAi experiments. The following strains have been used in this study: Bristol N2, *his-72(tm2066)*, *glp-1(e2141)*, *glp-4(bn2)*, MTP27(*gfp::3xflag::elo-6(S2A))*, MTP29 (*gfp::3xflag::elo-6(S2A);glp-1(e2141))*, MTP30 (*gfp::3xflag::elo-6(S2A);glp-4(bn2))*. MTP27 was generated by CRISPR-Cas9 system and further backcrossed with N2 for 3 times. MTP29 and MTP30 were generated by crossing MTP27 into *glp-1(e2141)* and *glp-4(bn2)* background. Metformin treatment was performed as previously reported ^33^. Metformin was added into agar at the concentration of 50 mM while preparing plates. For OP50 UV treatment, OP50 was seeded onto NGM plates and grew overnight at room temperature. The plates were then treated with 5 J/cm^2^ UV with the lids opened. Supplementation of C17iso was performed as previously reported ^39^. C17iso (Cayman) was dissolved in 100% DMSO at 10 mM concentration and mixed with OP50 to 1 mM final concentration before spotting plates, with equivalent DMSO concentration as the control treatment.

### RNA-seq library preparation and data analysis

Worm samples were collected, washed 3 times with cold M9, and frozen at -80 °C. Total RNA was extracted from frozen worm pellet using TRI reagent (Molecular Research Center). mRNA libraries were prepared and sequenced with PE150 at Annoroad for D2A to D12A aging samples or Novogene for GFP groups and *pqm-1* RNAi treatment. mRNA-seq data were analyzed as previously described ^21^. tRNA and rRNA reads were first filtered out using Bowtie, and the remaining reads were further aligned to WBcel235 by TopHat2(v2.1.1) with no novel junctions allowed ^40^. The uniquely aligned reads with a maximum of two mismatches were kept for differential expression analysis. The relative expression levels at different ages were calculated as FPKM using CuffDiff of Cufflinks software version 2.2.1^40^. Differential gene expression was analyzed by edgeR ^41^.

### CRISPR/Cas9 alleles generation

CRISPR/Cas9-based gene editing to knock in the *gfp::3xflag* coding sequence at the 5′-end of *elo-6* was performed as previously described ^42^. Repair templates in pDD282 (Addgene 66823) (forward primer for upstream arm: ACGTTGTAAAACGACGGCCAGTCGCCGGCATTGGCGCACAAATCACAAAA, reverse primer for upstream arm: TCCAGTGAACAATTCTTCTCCTTTACTCATTTTTACCTGCAATTTTAAACTTAAAAAAA, forward primer for downstream arm introducing S2A mutation: CGTGATTACAAGGATGACGATGACAAGAGAATGaCACAGGGAGAAGTCTCA, reverse primer for downstream arm: TCACACAGGAAACAGCTATGACCATGTTATTCACATCGCATTTCTGGCCC), sgRNA in pDD162 (CAGGGAGAAGTCTCATTCTTGTTTTAGAGCTAGAAATAGCAAGT), and a pharyngeal fluorescence selection marker pCFJ90 (Addgene 19327) were injected into N2 young adults. Homozygous knock-in alleles were verified by Sanger sequencing. The knock-in strains were backcrossed with N2 for at least 3 times.

### Lifespan experiments

For N2 background strains, embryos were laid and cultured at 20°C. For *glp-1(e2141)* and *glp-4(bn2)* background strains, embryos from 16°C cultured worms were laid at 16 °C onto lifespan plates, then hatched and were cultured at 25°C. Worms with different GFP levels were picked under the Leica M205FCA equipped with ET485/10x and 69000m filter for GFP ^43^. For RNAi treatment, worms were transferred to freshly IPTG induced iOP50 ^44^ or xu363 ^45^ RNAi plates at the specific age and further transferred every 1 or 2 days to freshly IPTG induced RNAi plates until approximately D10A. Lifespan was measured by scoring worms every other day or every day.

### RNAi treatment

All RNAi clone was obtained from the Ahringher bacteria library ^46^. Control L4440 and RNAi constructs were transformed into iOP50 or xu363. iOP50 or xu363 were grown in Luria broth with 50 µg/mL carbenicillin at 37°C overnight, concentrated 10 fold, and seeded on NGM plates. Bacteria were induced with 4 mM IPTG for 4 hours. Worms were transferred to freshly induced RNAi plates every day or every other day. For mRNA-seq of *pqm-1* RNAi treatment, MTP29 embryos from 16°C cultured worms were laid onto plates, hatched, and were cultured on OP50 until D2A at 25°C, then transferred onto the RNAi plates at 25°C and collected for RNA extraction on D5A.

### Body bending rate

Animals were individually transferred into S buffer (100 mM NaCl and 50 mM potassium phosphate, pH 6.0) and let rest for 1 minute. The numbers of body bending per 30 seconds in the liquid were counted single-blinded.

### Pharyngeal pumping rate

Pharyngeal pumping rates of animals were scored directly on NGM plates. The number of pharyngeal contractions was measured in 1 minute using a Leica M205FCA.

### Statistical analysis

Statistical analyses were performed by SPSS software or excel. The survival function was estimated using the Kaplan Meier estimator (SPSS software), and statistical analysis was done using the log-rank test or Breslow test. *P*-value < 0.05 was considered as significantly different from the control population, (∗∗∗) *P*-value < 0.001, (∗∗) *P*-value < 0.01, (∗) *P*-value < 0.05, (ns) *P*-value > 0.05.

Detailed description of tests performed to determine the statistical significance is included in figure legends.

### Motif analysis

Motif analysis of gene promoter region was performed by Homer (findMotifs.pl *genelist.txt* worm *output directory*/) using default setting (-start -400 -end 100 -len 8,10). Motif is annotated by Homer (findMotifs.pl *genelist.txt* worm *motif directory* -find *motif file > output*).

### Imaging

Images of worms on 3% agarose pad in 10 mM levamisole were taken with the Olympus BX53/DP74. Images of worms on NGM plates were taken with the Leica M205FCA/K3C. Both Olympus BX53 and Leica M205FCA were equipped with ET485/10x and 69000m filter for GFP imaging to avoid autofluorescence.

## Data availability

All data are available within this article and its Supplementary Information. The mRNA-sequencing datasets generated in this study have been deposited in the Genome Sequence Archive in National Genomics Data Center (NGDC) PRJCA021128.

## Acknowledgements

We thank the *Caenorhabditis elegans* Genetics Center for worm strains. We thank Dr. Leonard Krall for critical reading and comments. This work was supported by funding from the National Science Foundation of China (32160163) and Yunnan Fundamental Research Projects (202201BF070001-009) to M.P.

## Supplementary Table

**Table S1.** Genome-wide correlation analysis of mRNA-seq data

**Table S2.** Gene expression levels during aging

**Table S3.** Functional annotation of genes consistently decreased expression with age

**Table S4.** Differential gene expression between GFP groups

**Table S5.** Functional annotation of genes differentially expressed between GFP2 and GFP4 in both *glp-1(e2141)* and *glp-4(bn2)*

**Table S6.** Differential gene expression caused by *pqm-1* RNAi

**Table S7.** Phenotype summary of RNAi experiment targeting the genes highly expressed in the short-lived individuals

**Table S8.** Quantitative lifespan data

## Supplementary Figure

**Figure S1.**
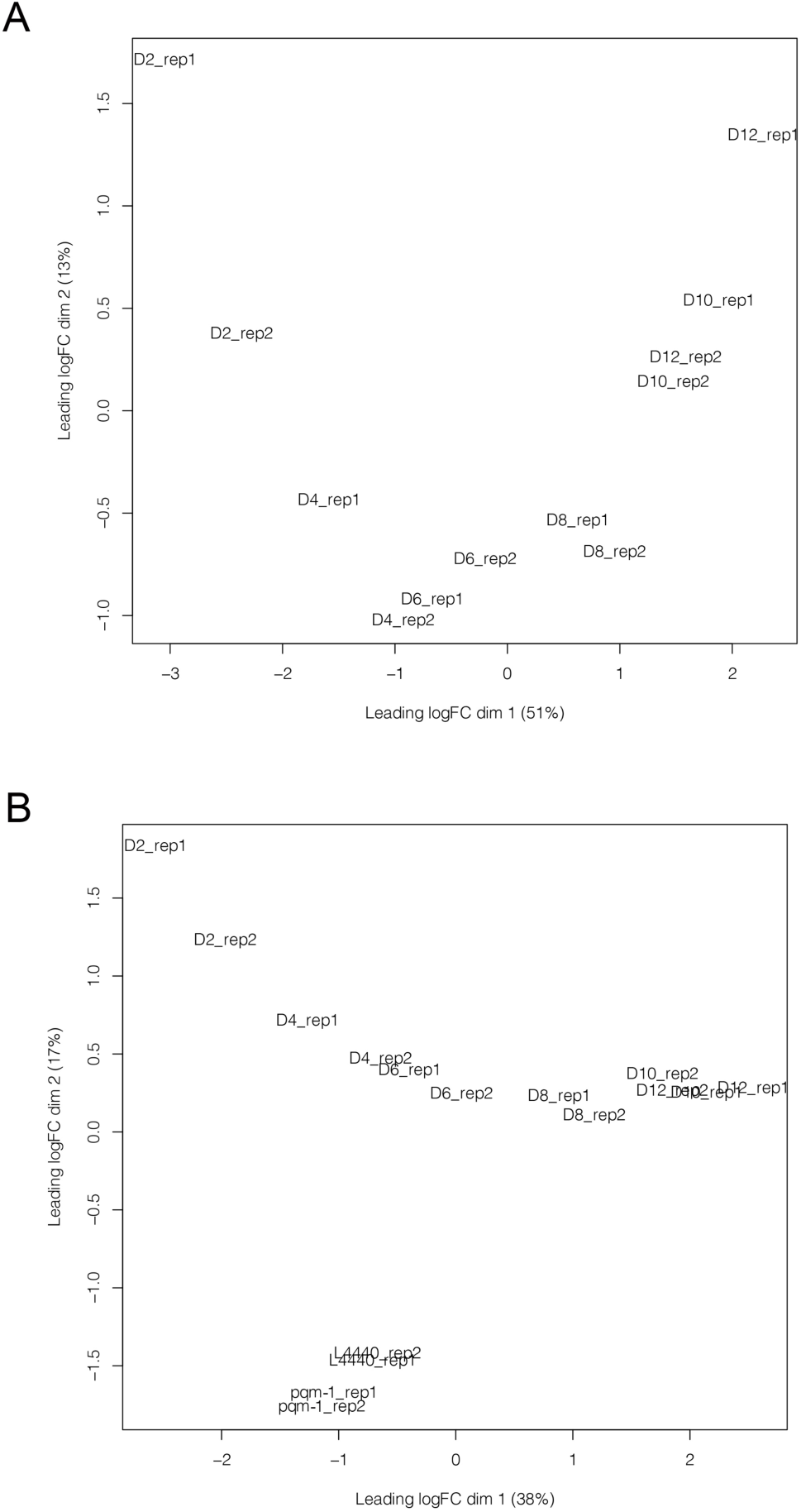
(A) MDS plot of mRNA-seq data from D2A to D12A in *glp-1(e2141)*. (B) MDS plot of mRNA-seq data in (A) and D5A in *glp-1(e2141)* with *pqm-1* RNAi treatment.

**Figure S2.**
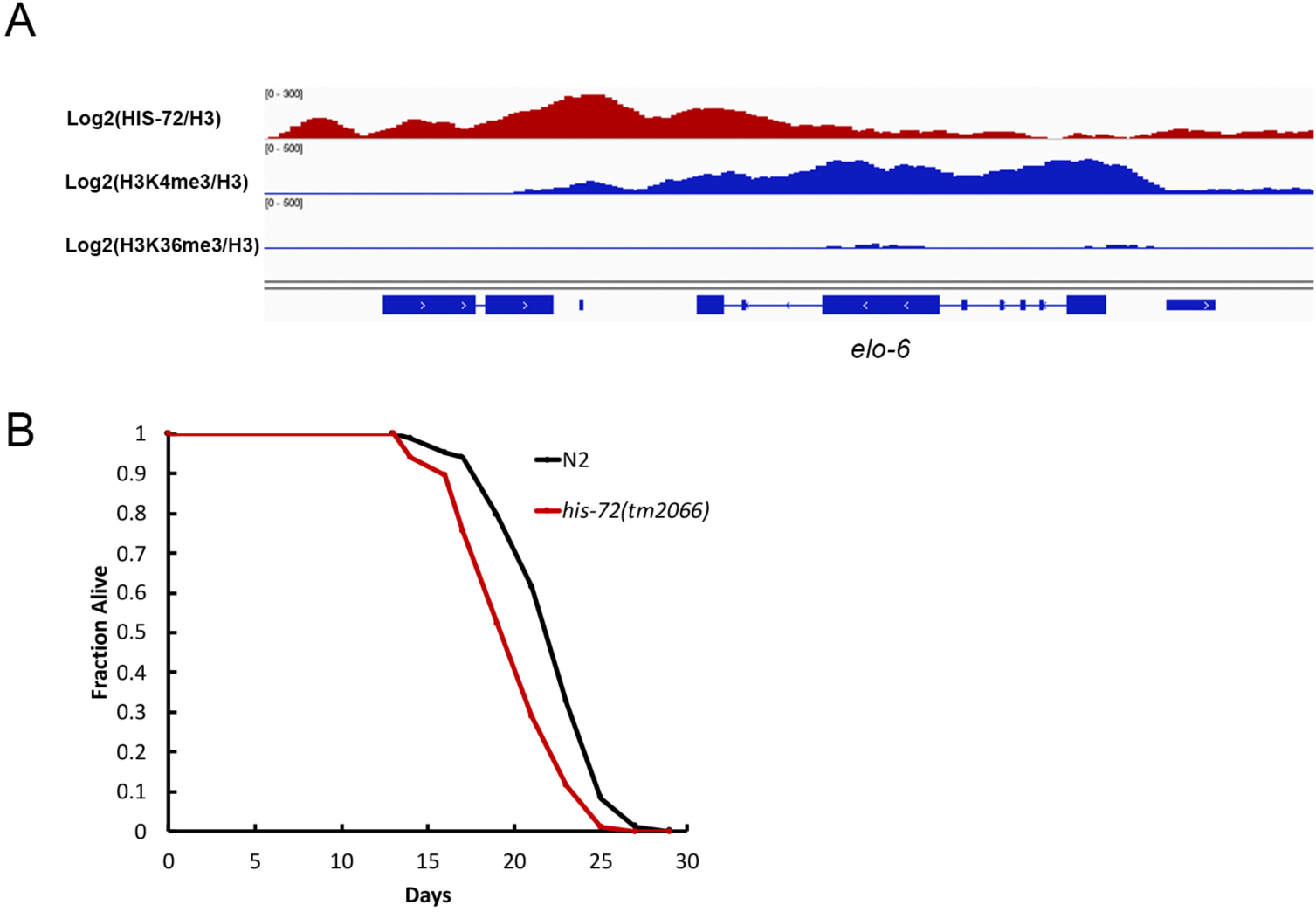
(A) The genomic region of *elo-6* is deposited with HIS-72, the adult-specific H3K4me3 on the gene body, and is absent of H3K36me3. (B) A loss-of-function mutation in *his-72* shortens lifespan. Quantitative data are presented in Supplementary Table S8. (∗∗∗) *P*-value < 0.001, *Log-rank test*

**Figure S3.**
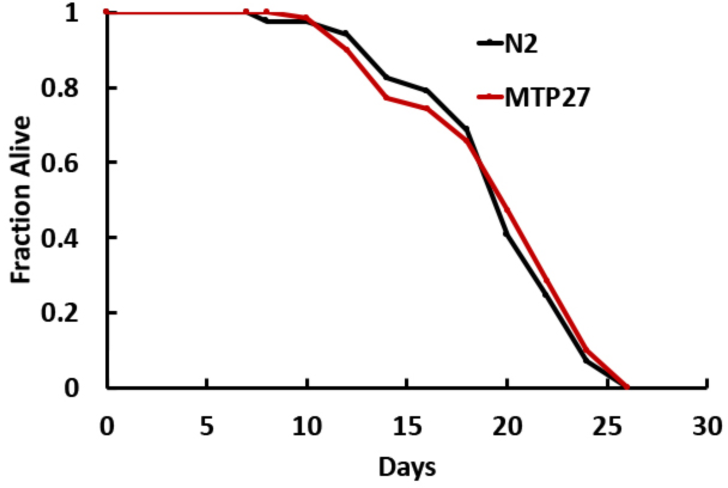
MTP27 has a similar lifespan as N2. Survival curves of N2 and MTP27 are shown. Quantitative data are presented in Supplementary Table S8. *P*-value > 0.05, *Log-rank test*

**Figure S4.**
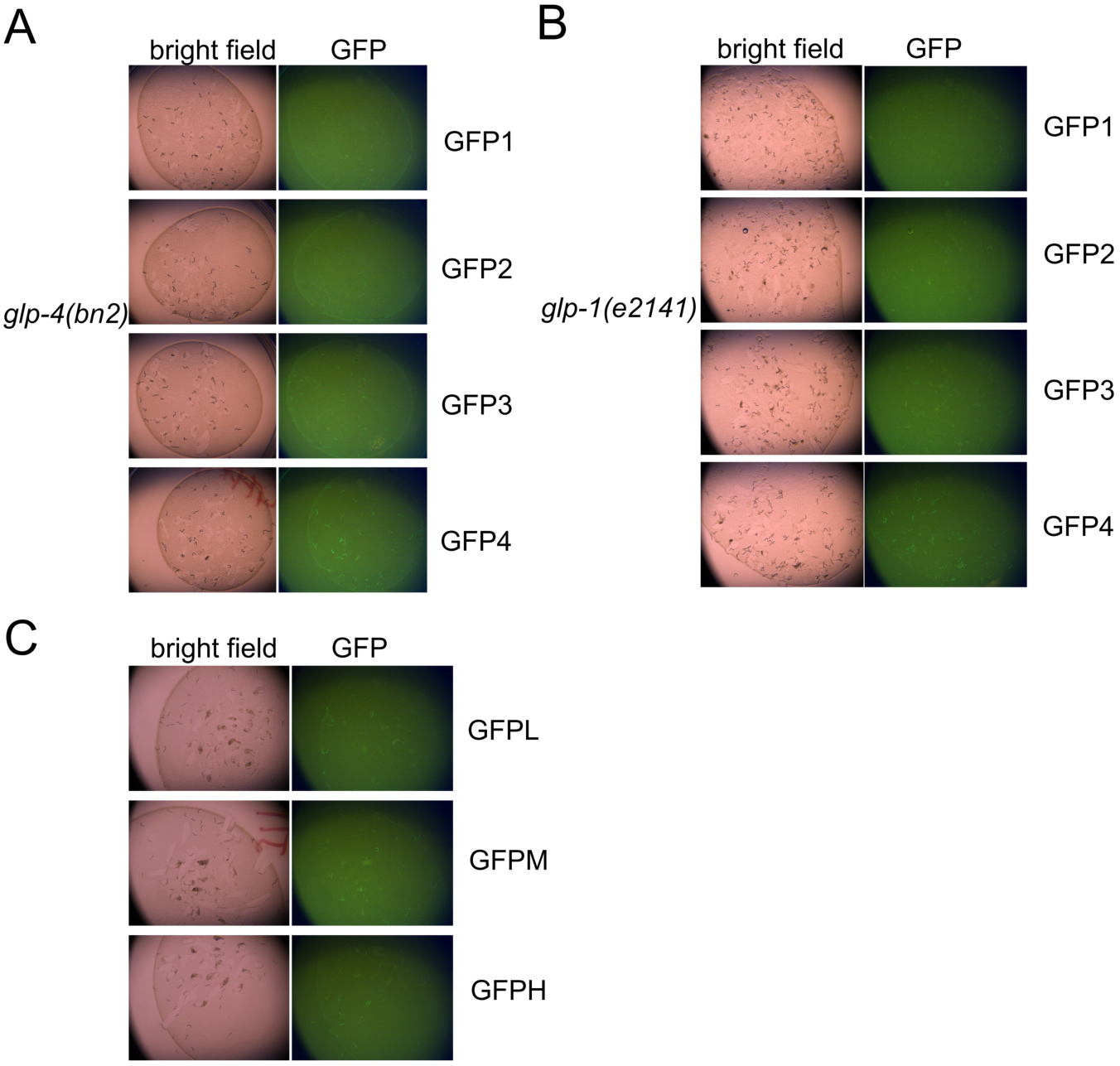
(A) The GFP expressions in GFP1, GFP2, GFP3, and GFP4 group worms from MTP30 on D5A. Images were taken by a fluorescence stereo microscope with consistent imaging parameter settings. (B) The GFP expressions in GFP1, GFP2, GFP3, and GFP4 group worms from MTP29 on D5A. Images were taken by a fluorescence stereo microscope with consistent imaging parameter settings. (C) The GFP expressions in offspring of GFPL, GFPM, and GFPH group worms from MTP27 on D5A. Images were taken by a fluorescence stereo microscope with consistent imaging parameter settings.

**Figure S5.**
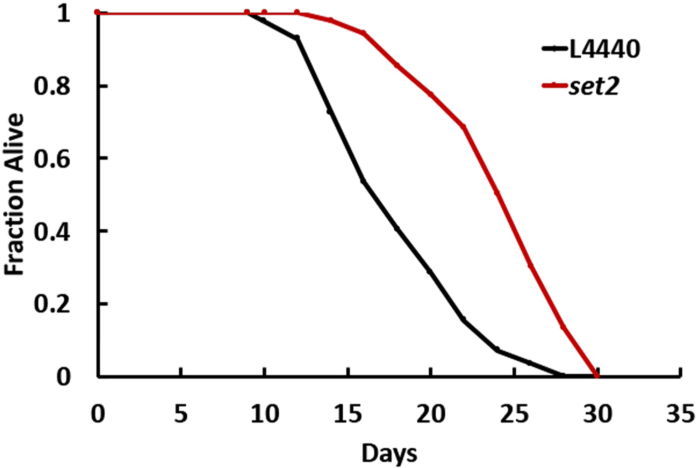
Survival curves of *set-2* RNAi and the L4440 control are shown. Quantitative data are presented in Supplementary Table S8. (∗∗∗) *P*-value < 0.001, *Log-rank test*

**Figure S6.**
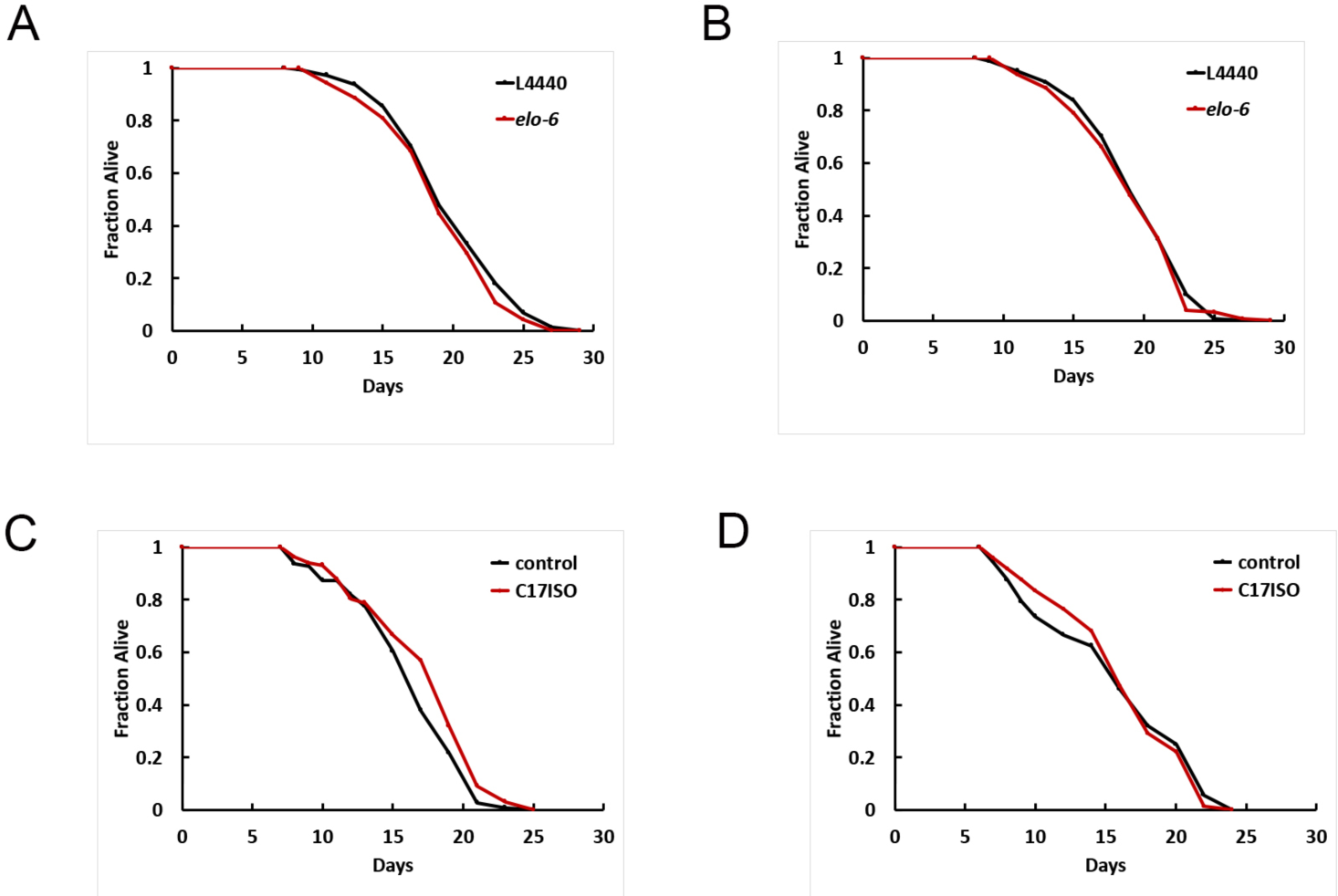
(A) *elo-6* RNAi treatment from D2A does not affect lifespan. Quantitative data are presented in Supplementary Table S8. (B) *elo-6* RNAi treatment from D5A does not affect lifespan. Quantitative data are presented in Supplementary Table S8. (C) Supplementation of C17iso from D4A slightly extends the populational lifespan, with a 3% mean lifespan extension compared to control. Quantitative data are presented in Supplementary Table S8. (∗) *P*-value < 0.05, *Log-rank test* (D) Supplementation of C17iso from D6A does not change the lifespan of GFPL group animals. Quantitative data are presented in Supplementary Table S8.

**Figure S7.**
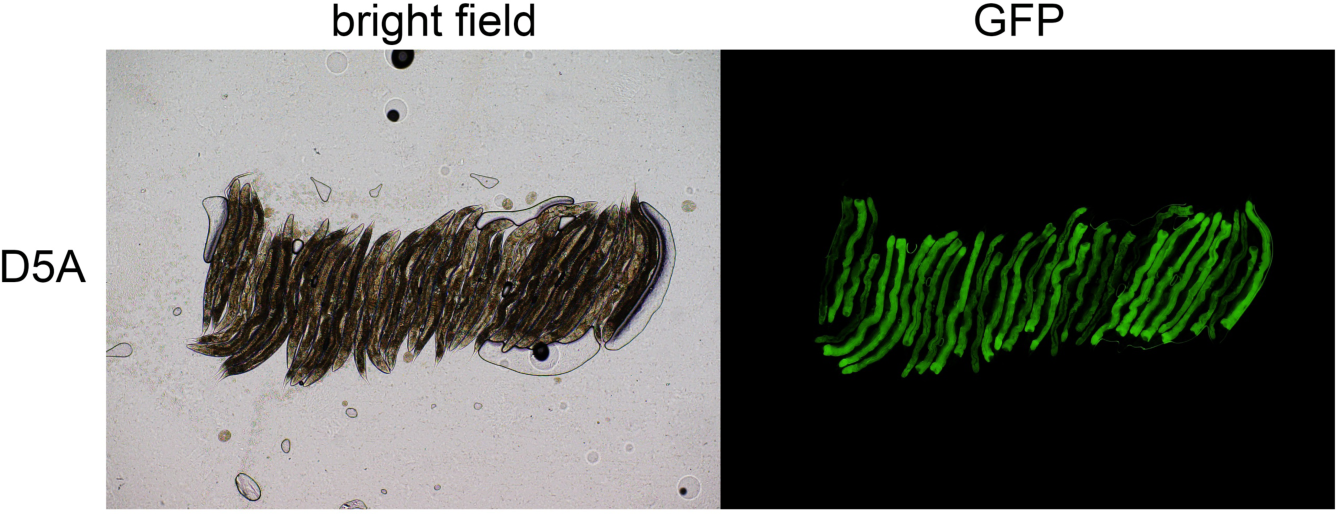
Expression of ELO-6 shows individual variation on adult day 5 when worms are fed with UV treated OP50 for 2 generations.

**Figure S8.**
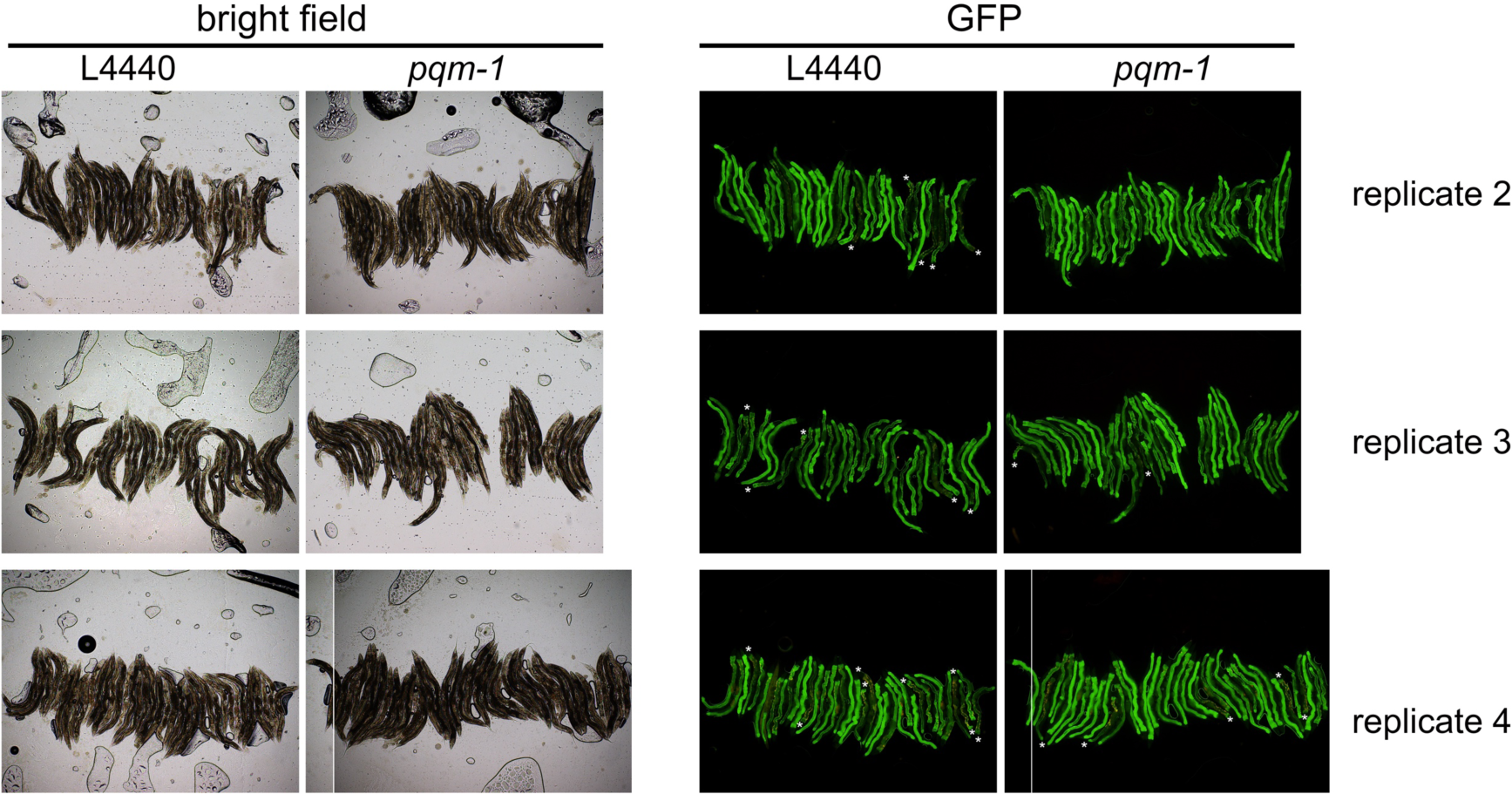
*pqm-1* RNAi treatment enhances GFP::ELO-6 expression homogeneity in MTP27 on D6A. RNAi treatment started from D3A. *: worms without GFP signal in part of the intestine.

**Figure S9.**
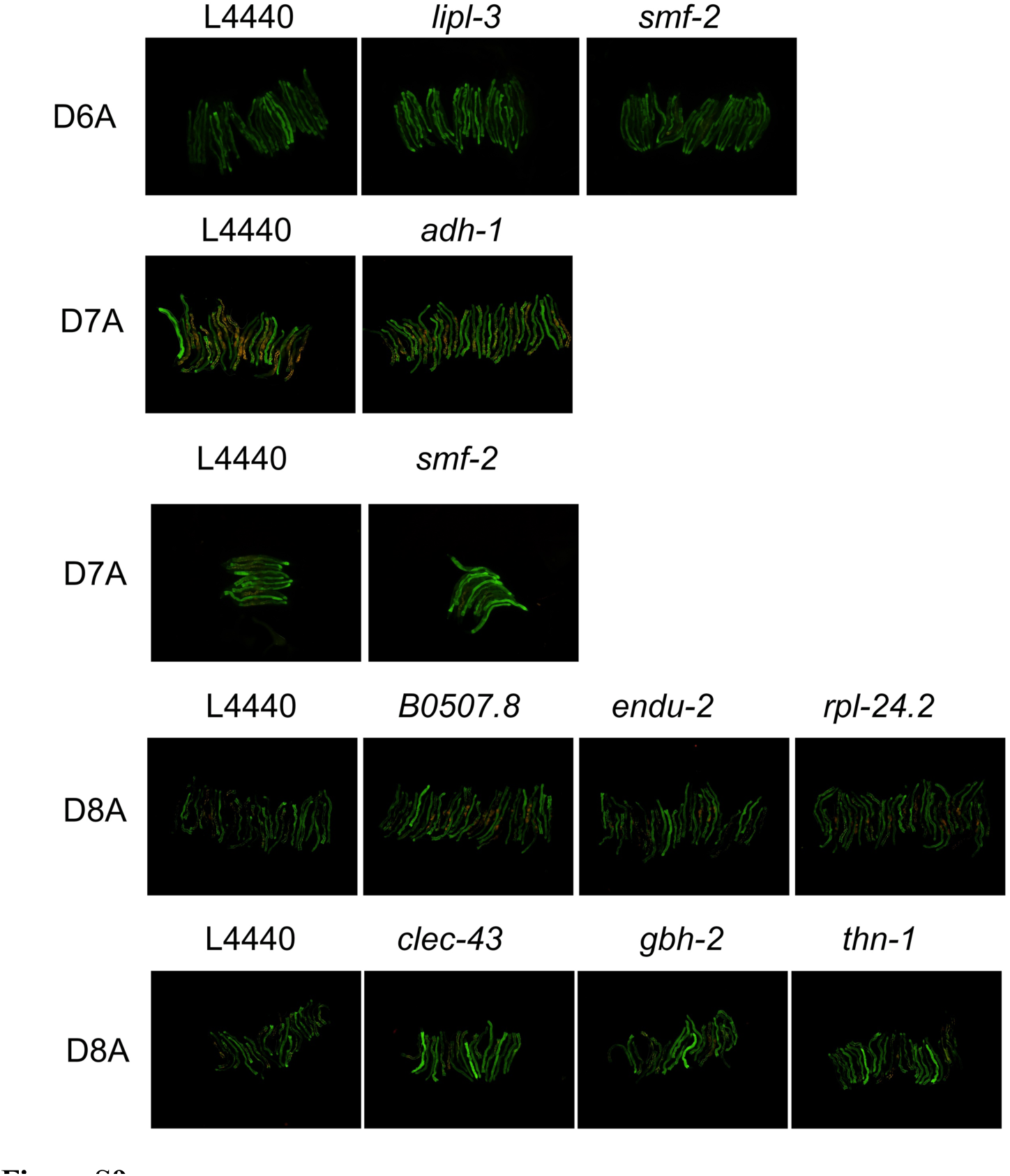
Representative images showing RNAi treatment of the genes highly expressed in short-lived individuals from adult day 2 enhances ELO-6 expression stability with age.

